# P-BODY AND CYTOSKELETON REMODELING BY ORTHOHANTAVIRUSES

**DOI:** 10.1101/2024.07.16.603662

**Authors:** Hannah Sabeth Schwarzer-Sperber, Annette Petrich, Matthias Schade, Niklaas Nilson, Linah Chibrac-Ahad, Maik Joerg Lehmann, Katharina Paulick, Sabrina Weiss, Daniel Bourquain, Peter T. Witkowski, Detlev H. Krüger, Andreas Herrmann, Roland Schwarzer

## Abstract

Orthohantaviruses, are emerging zoonotic pathogens causing life-threatening diseases in humans. The orthohantavirus genome consists of three RNA segment (vRNAs) of negative polarity, which are encapsidated by the viral nucleoprotein (N). To date, the precise subcellular behavior of vRNAs and N has not been fully elucidated. Here, we present a comprehensive analysis of orthohantavirus infections using Fluorescence *in situ* Hybridization (FISH) and multiple sequential FISH (Mu-Seq FISH), which enables simultaneous detection of viral RNAs, viral mRNAs, N, and cellular factors. Our experiments revealed distinct patterns of viral RNA clustering with varying degrees of N association. Moreover, we found a significant spatial correlation of virus vRNAs and N with cellular processing (P)-bodies, underlining their key role in orthohantavirus replication. Throughout the course of an infection, we observed an increasing dominance of N expression, while concomitantly P-body numbers grew significantly. We also found indications for a preferential 5’-end degradation of viral mRNAs in P-bodies. Furthermore, we report that orthohantavirus infection is accompanied by a significant redistribution of cellular components: while filamentous actin and microtubules become enriched in the perinuclear region, P-bodies move to the cell periphery. Finally, co-localization analyses suggest a formation of viral factories containing N, vRNAs, and viral mRNAs, indicating an intricate orthohantavirus assembly hierarchy.

**AUTHOR SUMMARY:** In this study we used advanced imaging techniques to observe the dynamics of viral components and key cellular structures during orthohantavirus infections. Our experiments show that orthohantaviruses cause significant changes within the cell, particularly involving P-bodies and components of the cytoskeleton, such as actin and microtubules. We also provided comprehensive spatiotemporal maps of orthohantaviral components, including visualization of the viral nucleoprotein, genomic RNA and mRNAs. Finally, we found indications for a 5’end degradation of virus mRNA in P-bodies, thus adding to our understanding of intracellular host-pathogen crosstalk. In summary, our work highlights the intricate relationship between viruses and host cells, emphasizing the dynamic changes that occur during orthohantaviral infections.

## INTRODUCTION

Orthohantaviruses (family *Hantaviridae*) are a group of emerging, globally distributed viral pathogens that can cause serious and sometimes fatal diseases in humans (1). Orthohantaviruses harbor a tri-segmented, negative sensed, single-stranded RNA genome with viral genomic RNA (vRNA) segments designated S- (small), M- (medium) and L (large). The S-vRNA encodes for the viral nucleoprotein N, whereas M and L contain ORFs of the glycoprotein precursor GPC and the RNA-dependent RNA polymerase (RdRP), respectively. Some Orthohantaviruses harbor an alternative reading frame in the S-vRNA that encodes a non-structural protein (NSs) with accessory functions (1).

To date, the exact site the of the orthohantavirus replication has not been identified (2), however cellular processing bodies (P-bodies), RNA storage and RNA degradation sites, have been reported to play a central role (2–4). Of note, orthohantavirus infections have been suggested to involve the formation of virus factories (2,5) as previously described for the closely related Bunyamwera virus species (6), however clear evidence for such entities in orthohantavirus infected cells is still missing. Despite having a comprehensive understanding of the fundamental mechanisms and key processes underlying orthohantavirus lifecycles in the host cell, there remains a scarcity of knowledge concerning their spatiotemporal dynamics. Important questions about the complex crosstalk between different virus components, as well as between pathogen and host, remain unanswered to this day.

Here, we used Fluorescence *in situ* hybridization (FISH) and immunofluorescence staining (IF) to investigate the localization and dynamics of all three orthohantavirus vRNA segments, virus mRNAs, and the viral N protein by fluorescence microscopy. Moreover, we analyzed key cellular factors, majorly contributing to orthohantavirus entry, replication and assembly, namely actin, tubulin and P-bodies.

## MATERIALS AND METHODS

### Cell culture and infection

PUUV infection (strain Sotkamo: V-2969/81 Charite University Hospital, Berlin, Germany) were conducted in African green monkey kidney epithelial cells (Vero E6, ATCC CRL- 1586; American Type Culture Collection, Manassas, VA) maintained in Dulbecco’s modified Eagle medium (DMEM) supplemented with 10% heat-inactivated fetal bovine serum (FBS), 2 mM L-glutamine, 100 U/ml penicillin, and 100 μg/ml streptomycin (all from PAA Laboratories GmbH, Austria) under standard cell culture conditions. Cells were passaged every 2-3 days when they reached nearly 80 % confluence in tissue culture flask, for no more than 15 cycles.

PUUV infection was performed as previously described (7). Briefly, cells were grown to 50-60% confluency, rinsed once with Dulbecco’s phosphate-buffered saline with Mg^2+^/Ca^2+^ (DPBS+/+) and infected with a multiplicity of infection (MOI) of 0.3 - 1 in DMEM containing 1% FBS. After an incubation of one hour at 37°C, cells were washed three times with DPBS+/+ to remove unbound viruses. Cells were further incubated with DMEM supplemented with 1% FBS at 37°C. Fixation, staining and microscopy was performed at 72 hpi if not otherwise stated, to ensure robust hantavirus infection levels and advanced replication stages (8). All solutions, buffers, and media used for cell culture were purchased from PAN-Biotech (Aidenbach, Germany).

### Immunofluorescence and DAPI staining

Cells were incubated with 0.2% acetylated bovine serum albumin (BSA) (B8894, Sigma- Aldrich, St. Louis, MS, USA) in 2× SSC buffer supplemented with 2 mM of the unspecific RNase inhibitor vanadyl ribonucleoside complex (VRC, Sigma-Aldrich, St. Louis, MS, USA) for 15 min at RT, followed by incubation with a cross-reactive anti-Tula virus/Malacky N protein antibody (1:500) (9), anti-Dcp1a for P-body detection (1:100, #ab57654, Abcam, Cambridge, UK), anti-α-tubulin (1:1000, #T5168, Sigma-Aldrich, St. Louis, MS, USA) and secondary antibodies, conjugated with different AlexaFluor 488 or 647 (1:1000, Thermo Fisher Scientific, Waltham, MA, USA) at dilutions of 1:100 to 1:1,000 in BSA-containing 2× SSC buffer for 45 min, again at RT. Samples were washed twice to remove unbound antibodies with 2× SSC buffer for 10 min at RT. Subsequently, cells were stained with either 100 nM 4′,6-diamidino-2-phenylindole (DAPI) (Thermo Fisher Scientific, Waltham, MA, USA) or Hoechst 33342 (Thermo Fisher Scientific, Waltham, MA, USA) at RT for 10 min to label DNA. Finally, the samples were again washed twice with 2× SSC. Filamentous actin was stained with TRITC conjugated phalloidin (#P1951, Sigma-Aldrich, St. Louis, MS, USA) according to the manufacturer’s protocol.

### Fluorescence *in situ* hybridization

FISH probe design is described in the supplementary methods (SM). In a typical FISH experiment, samples were first treated with pre-warmed 80% formamide in 2×SSC at 37 °C for 10 min, to enhance FISH staining signals by increasing the accessibility of vRNA for probes. Subsequently, samples were rehydrated with 2× SSC buffer at RT for 10 min. Then, cells were incubated with hybridization buffer (200 nM FISH probes, 2× SSC, 10% formamide, 10% dextrane sulfate and 2 mM VRC) at 37 °C for 2–4 h, followed by two washing steps with pre-warmed 10% formamide in 2× SSC at 37 °C for 10 min. For FISH probes removal between sequential labelling steps (multiple sequential-FISH, MuSeq-FISH) an 80% formamide buffer wash was used, which decrease the melting temperature of double-stranded nucleic acids (10). Of note, treatment with high concentrated formamide buffer impairs immunofluorescence staining, so all MuSeq-FISH experiments were conducted after antibody staining. In a typical MuSeq-FISH experiment, samples were imaged and then subjected to oligonucleotide probe removal by washes with pre-warmed 80% formamide at 37 °C for 10 – 15 min followed by rehydration with 2× SSC at RT for 5 min. Prior to the initiation of new FISH cycle, successful removal of probes was verified by microscopy. Each FISH staining cycle labelled two different target RNAs and nuclei simultaneously. Last run of the MuSeq-FISH: HCS CellMask deep red (Thermo Fisher Scientific, Waltham, MA, USA) staining according to the manufacturer’s protocol; in combination with IF/DAPI staining.

### Microscopy and image analysis

Stained samples were subjected to fluorescence microscopy, followed by manual or automated, quantitative image analysis. Details can be found in the supplementary methods.

### Creation of hierarchical assembly trees

A regression-based model for the linear relationship between the abundances of multi-segment complexes (MCC) of a certain complexity (rank *k, k ∊* [2,7]), and the products of the abundances of their constituent components with the corresponding rank *k*–1 was developed and thoroughly tested previously (10) (available via GIT-HUB (https://github.com/Budding-virus/Packbund). As an example, an MCC composed of the components Cx, Cy, Cz could have formed in three ways: 1) Cx joined a MCC composed of Cy and Cz, 2) Cy joined a MCC consisting of Cx and Cz, or 3) Cz joined the MCC composed of Cx and Cy. The most likely assembly order is then estimated by taking into account the abundance of Cx, Cy, Cz and the respective binary complexes (*k*-1). The complete assembly tree for larger complexes is formed iteratively.

### Statistical analysis

If not stated otherwise, bars charts show arithmetic mean ± SEM. Statistical significance was assessed using GraphPad Prism version 10 (GraphPad Software Inc., San Diego, CA, USA), applying parametric one-way analysis of variance (ANOVA) and Tukey’s multiple comparisons tests and displayed as follows: **** p < 0.0001; *** p < 0.001; ** p = 0.001–0.01; * p = 0.01–0.05. All data were tested for normality by Shapiro–Wilk test using a significance level of 0.05.

## RESULTS

To systematically investigate orthohantavirus infection by FISH we have systematically designed probes for all three genomic viral RNA segments as well as for the respective virus protein coding messenger RNA sequences. First, we focused on PUUV vRNAs and tested whether we could reliably detect and visualize these viral nucleic acids and proteins concomitantly.

### FISH reveals orthohantavirus vRNA clusters with different levels of N protein association

For a proof-of-concept, we performed FISH and IF at 72 hours post infection (hpi), a rather late time point where robust infection levels are to be expected. Our experiment showed virus specific FISH signals for all three segments in infected, but not uninfected cells (Figure 1A, Figure S1A, B). Moreover, we observed distinctive populations of intracellular spots with only vRNA, only N protein or both (Figure 1A, Figure S1C).

**Figure 1:**
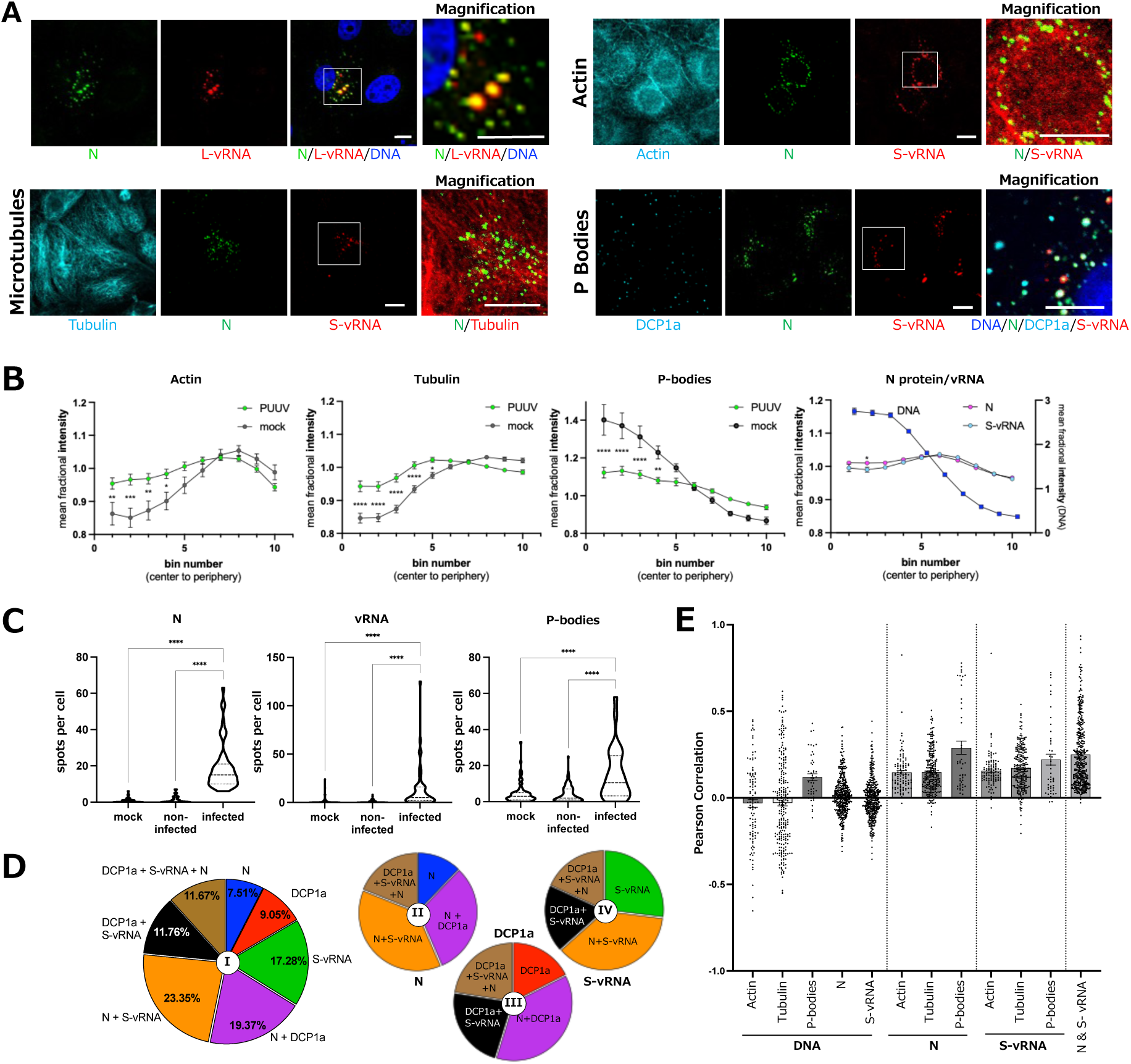
Immunofluorescence and FISH analysis of PUUV infected cells at 72 hpi. **(A)** Confocal images of PUUV infected (MOI=1) VeroE6 cells, subjected to IF staining and vRNA FISH at 72 hpi. The upper left panel shows N co-localization with vRNA, while all other panels focus on co-localization of viral molecules with cellular markers. Colored labels indicate the displayed target molecule. The boxed area in the overlay image is shown magnified on the right. The upper left panel shows maximum intensity projections of z-stacks, obtained by confocal microscopy. All other panels show equatorial slices. Hoechst 33342 or DAPI were used as a DNA counter staining (blue). Scale bars = 10 µm. (B) Automated image quantification was conducted using CellProfiler. Mean fractional intensity (MfrI) was calculated using the CellProfiler module MeasureObjectIntensityDistribution, binning the cell body into 10 sections starting from the center of the nucleus going outwards. Several dozen to several hundred cells from different fields of view were analyzed for each plot. Error bars show the standard error of the mean (SEM), and statistical significance was assessed by comparing PUUV and mock samples at each bin. The far-right plot indicates an analysis of N protein and S-vRNA intensities in infected cells. DNA MfrI is included as control. (C) Numbers of identified N protein, P-body and S- vRNA spots in untreated (mock), virus exposed but non-infected, and virus infected cells displayed in violine plots. Infected and non-infected cells were distinguished based on the N protein signal. Experiments were repeated three times; n = 847 cells. ****, p≤ 0.0001. Significance was analyzed using two-way ANOVA and Tukey’s multiple comparisons test. (D) Co-localization of identified objects. Each section of the pie charts shows the percentage of a specific spot population with respect to all other spots in the respective analyses. The pie chart I summarizes all identified spot species (N, S-vRNA and DCP1a), whereas the smaller charts only focus on spots positive for N (II), DCP1a (III), or S-vRNA (IV), respectively. (E) Pearson correlation calculated per cell between indicated stainings. Individual cell measurements are shown as single dots, and all bars indicate mean with SEM. Statistics are provided in SI table S1.

### Viral RNA associates with actin accumulations, microtubules and P-bodies

In the next step we sought to assess if and to what extent vRNAs spatially correlate with different cellular factors. We focused on three key host components that have previously been described, by us and others, to play important roles during orthohantavirus infection: actin, microtubules and P-bodies (3,6,7,11,12). Importantly, in our previous studies we have shown that infected cells exhibit a significant enrichment of filamentous actin in the perinuclear region when compared to mock-infected cells (7). Here, we found indications that both N protein and PUUV vRNAs co-localize with these actin accumulations (Figure 1A, Figure S2A, C). Similarly, N and vRNAs appeared to be lined along cellular microtubules (Figure 1A, Figure S2A, C). We further detected a considerable co-localization of all three vRNA segments and host P-bodies (Figure 1A, Figure S2D-F), which was in many cases accompanied by N protein positivity in P-body/vRNA double-positive spots.

### PUUV infection induces remodeling of actin filaments, microtubules and P-bodies

To enable an unbiased and high-throughput, quantitative examination of our microscopy images, we next turned to automated image analysis of large data sets that we acquired across multiple experiments. For that purpose, micrographs were thoroughly analyzed using the CellProfiler software with an in-house customized processing pipeline, designed to segment, classify and quantify images at the single-cell level (Figure S4). Briefly, our pipeline identifies nuclei based on DNA counterstaining, followed by recognition of cellular bodies in either IF stainings of abundant cellular markers (such as microtubules or actin), or in transmission light images (differential interference or phase contrast). Then, P-Body, N protein or vRNA puncta were identified independently. Finally, identified objects were saved as regions of interest (ROI) and quantitatively assessed.

First, we asked whether the overall distribution of the three cellular proteins, actin (actin filaments), α-tubulin (microtubules) and DCP1a (P-bodies) changed upon infection. To this aim, we calculated the mean fractional intensity (MFrI) of the respective proteins as a function of their distance to the cell center and compared infected vs. mock-infected cells. Importantly, actin, α- tubulin and P-bodies were found to exhibit an aberrant intracellular distribution in infected cells (Figure 1B), specifically in the center and perinuclear region (bin 1-5). Whereas actin and tubulin appeared to significantly enrich in the cell center, P-bodies redistributed from the inner to the outer regions of the cells (Figure 1B). DNA was not displaced as a result of orthohantavirus infection (Figure S5). For comparison, we also assessed the distribution of N protein and vRNAs in infected cells. We focused on the S-vRNA, since this segment encodes for the viral N protein, thus allowing us to monitor the viral gene and its respective protein concomitantly. N protein on the other hand, was chosen due to its crucial role and high abundance in orthohantavirus infected cells. Of note, in contrast to the three investigated cellular proteins, viral RNA and N protein showed markedly less divergent signals in the central vs. the peripheral regions of the cells (Figure 1B, MFrI around 1) and hardly any statistical significance in their overall MFrI distribution.

### P-body numbers are increased in PUUV infected cells

Next, we quantified the overall number of identified N protein, DCP1a (P-bodies) and S- vRNA spots per cell for three different categories of cells: 1) PUUV infected, 2) non-infected cells in PUUV exposed cell cultures, and 3) mock infected (virus naïve) samples (Figure 1C). For that purpose, in PUUV-exposed samples, cells were classified into infected and non-infected cells based on a N protein signal threshold that was defined in mock infected samples. Naturally, this resulted in a strong significant difference in the number of N protein spots between mock or non-infected cells when compared with infected cells. On average, between 10 and 20 N protein spots were identified in infected cells whereas false positive spots were very rare (Figure 1C). An average number of around 13 S-vRNA spots was found in N protein positive cells, whereas N protein negative cells, again, barely showed any false-positive spots, indicating that our staining and analysis pipeline enables reliable and consistent identification of different viral molecules in infected cells. Interestingly, we observed a highly significant increase in the number of P-body spots in infected over both, non-infected cells and mock samples, which again points to a remodeling of cellular structures upon PUUV infection (Figure 1C). Importantly, we also found a significant, albeit less extensive redistribution and remodeling of both actin and P-bodies in a primary cell system based on human pulmonary microvascular endothelial cells (Figure S6), which indicates that this process is not cell-type specific.

### Extensive co-localization of P-bodies with N and S-vRNA

We then quantitatively evaluated co-localization of P-bodies and viral components in infected cells. We found that almost half of all identified spots (either S-vRNA, N or DCP1a positive) exhibited the P-body marker and at least one of the viral molecules (Figure 1D, pie chart I: brown, magenta and black sections). Focusing on P-bodies it became apparent that the large majority of all DCP1a positive spots were associated with either N, S-vRNA or both (Figure 1D, pie chart III). Similarly, N protein and S-vRNA were largely co-localizing with P-bodies or each other, and only a small fraction was found non-associated with any other investigated factor (Figure 1D, pie charts II and IV, blue and green sections). Overall, these findings demonstrate a substantial spatial correlation of N protein, S-vRNA and cellular P-bodies in PUUV infected cells.

Lastly, we assessed the Pearson correlation of all staining, by investigating pixel-by-pixel co-localization of the target molecules under study (Figure 1E). Of note, this analysis does not require identification of spots, nor are thresholds or other user-defined parameters necessary, so that obtained results are particularly un-biased and objective. In agreement with our above- described findings, we observed the highest, positive correlation between N proteins and P-bodies, S-vRNA and P-bodies, as well as N protein and S-vRNA (Figure 1E). A more moderate positive correlation of N protein and S-vRNA was found with actin and α-tubulin, but interestingly also between DNA and P-bodies, likely indicating a nucleus-adjacent localization of P-bodies in PUUV-infected cells (Figure 1E). As expected, no correlation was found between DNA and actin, α-tubulin, N protein or S-vRNA (Figure 1E).

### Using MuSeq-FISH for parallel imaging of vRNAs, mRNAs and N protein

The orthohantavirus life cycle involves three key types of viral molecules: the vRNA, the corresponding mRNA and, ultimately, the encoded viral protein. To take this into account, we designed a novel set of FISH probes for the parallel detection of vRNA, mRNA and N protein, thus covering the entire molecular replication sequence of the PUUV S segment (S-vRNA, S- mRNA and N). Moreover, in order to be able to visualize all viral factors simultaneously, we employed a recently developed technique that enables the detection of numoerus nucleic acid species in a single FISH experiment (10), called Multiple Sequential FISH (MuSeq-FISH). The principle behind this method (see also Figure S7) is that RNAs are detected with a first set of FISH probes, followed by a stringent formamide wash that removes all bound FISH probes (comparable to membrane stripping in immunohistochemistry). Subsequently, another round of FISH staining can be conducted, with a different set of FISH probes. Since washing and staining can be repeated multiple times, several nucleic acids can be detected and discriminated in the same fluorescence channels. Here, we applied this approach to stain PUUV infected cells for all viral nucleic acid species (S-m/vRNA, M-m/vRNA, L-m/vRNA,) as well as N protein and P-bodies over extended periods of time (up to 240 h). In addition, we used DAPI as a DNA counterstain and CellMask staining to facilitate subsequent, automated image segmentation and identification of cell bodies. Representative micrographs at 240 hpi are shown (Figure 2, Figure S7). Importantly, we again found numerous spots with only single stainings (Figure 2B), which indicates a high specificity of all utilized FISH probe sets.

**Figure 2:**
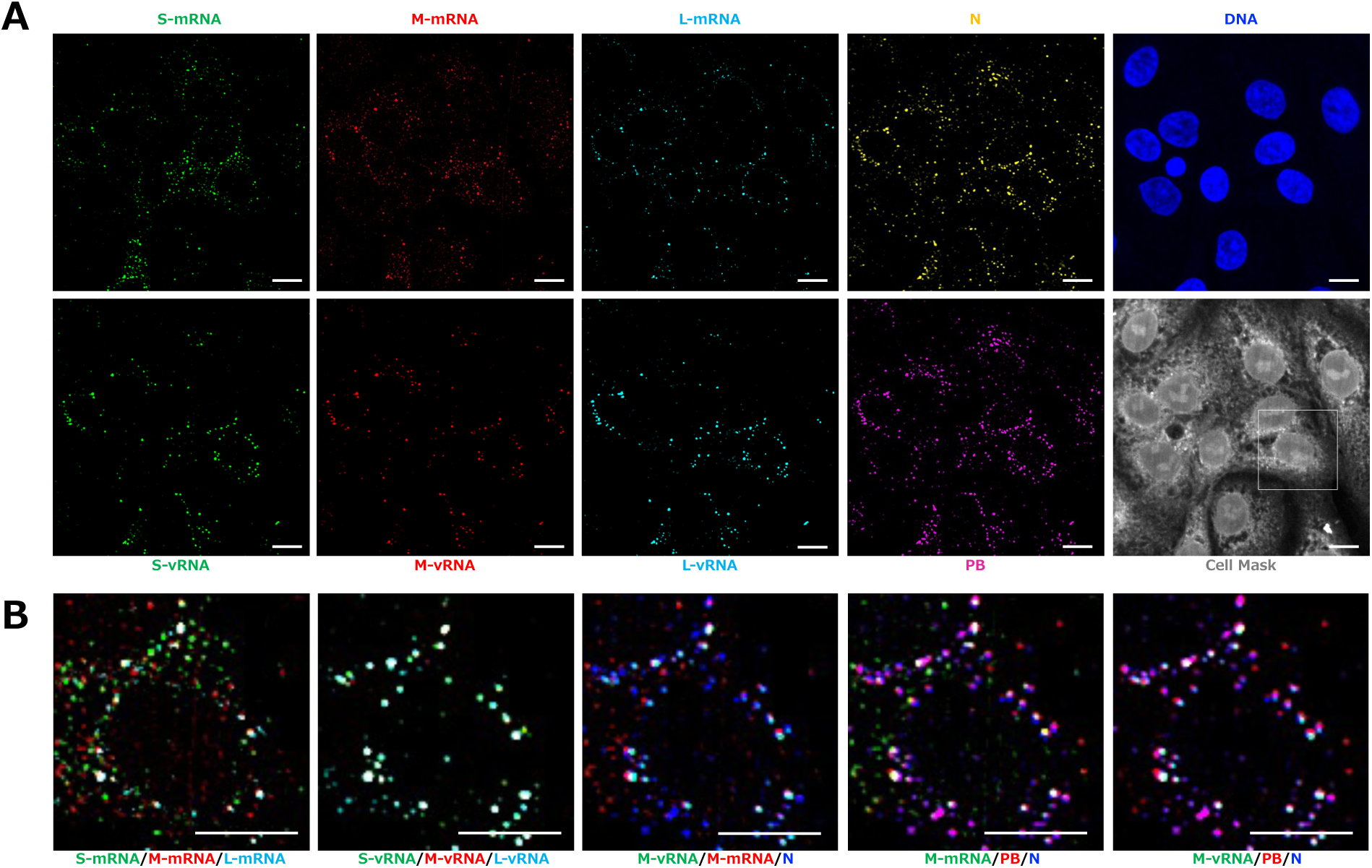
Utilization of Multiple sequential FISH (MuSeq-FISH) to image PUUV vRNAs and their transcripts in infected cells. VeroE6 cells were infected with PUUV (MOI=0.3) for 240 h and analyzed using MuSeq-FISH, enabling consecutive imaging of multiple, viral nucleic acid species, viral and cellular proteins and DNA. **(A)** Overview images of a representative region of interest, showing each staining individually: S-mRNA and S-vRNA in green, M-mRNA and M-vRNA in red, L-mRNA and L- vRNA in cyan, N protein in yellow, P-bodies (PB) in purple, DNA in blue and CellMask staining in grey. **(B)** Magnification of the boxed region shown in the cell mask image in (A). Overlay images are shown as indicated below. Scale bars=10 µm. Images represent maximum intensity z projections. Additional images can be found in Figure S8.

### Quantitative analysis of MuSeq-FISH shows increasing abundance of viral mRNAs, vRNAs, N protein and P-bodies throughout orthohantavirus replication cycles

The comprehensive Mu-Seq-FISH data sets were investigated using FISH-quant, a publicly available software for spot detection in 3D microscopy images (13), and with a custom-written R script for localization and co-localization analysis (10). In contrast to our previously used Cell Profiler pipeline, this approach detects and measures vRNA, mRNAs, N protein and P-bodies in three dimensions, which improves and refines image segmentation, and should result in overall higher numbers of identified objects.

Our MuSeq-FISH stainings at 24, 48, 72, 120, 168 and 240 hours post infection revealed a significant and gradual increase in the abundance of N protein, vRNAs, viral mRNAs and notably again of P-bodies (Figure 3A). We also quantified the distance of P-bodies and N protein puncta from the cell nucleus (Figure 3B), which was found to change significantly between the different time points. For P-bodies the mean distance did not follow a clear trend, but N protein was found in increasing distances from the nucleus over time, with a maximum at 168 h. This could be indicative of heightened virus trafficking to the cell surface at late stages of the viral replication cycle. Next, we investigated individual vRNA and viral mRNA species independently, again assessing overall abundance and distance to the cell center as a function of the infection time (Figure 3C). Expectedly, all three vRNAs and their respective transcripts were found in gradually increasing numbers, as the infection spread and progressed in the cell culture (Figure 3C). Interestingly, the intracellular distribution significantly diverged across the different mRNA species and even more pronounced between viral mRNA and vRNA (Figure 3C). Whereas S- mRNA and M-mRNA distance to the cell center significantly increased over time, L-mRNA showed a trend towards shorter distances, albeit not significantly. In contrast, all vRNAs showed a high distance peak at 24 hpi, followed by a significant drop at 48 h and again a gradual increase at later infection time points. We suspect the early peak to be reflective of the virus inoculum and thus convoluted by virus entry, while the following gradual increase might indicate true peripheral egress events.

**Figure 3:**
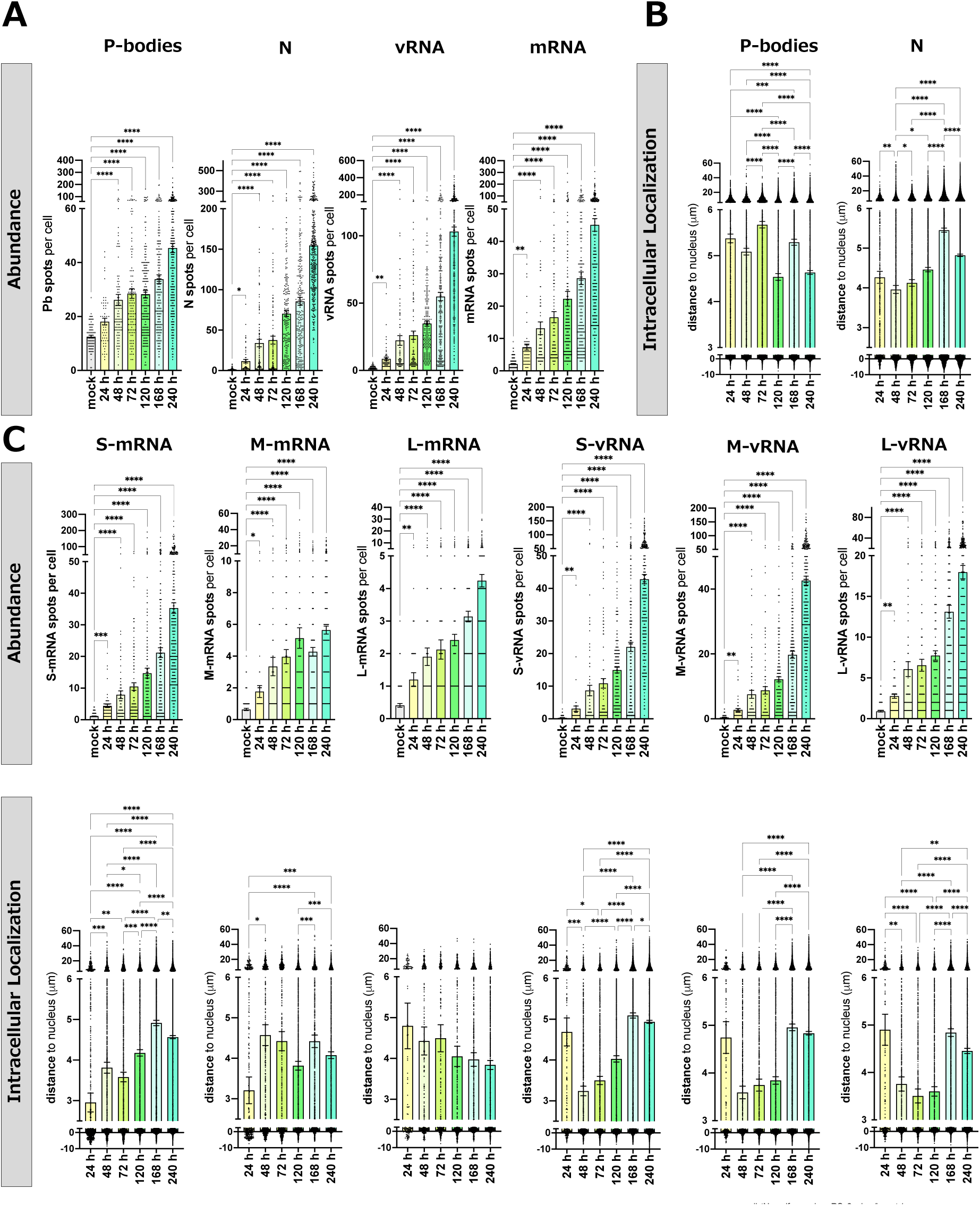
Quantitative analysis of PUUV infection kinetics using MuSeq-FISH over 10 days of infection. VeroE6 cells were infected with PUUV and investigated at different time points post infection using MuSeq-FISH. **(A)** Per cell counts of P-body, N protein, all vRNA species and all viral mRNA speckles in non-infected and infected cells. **(B)** Intracellular localization analysis, assessing the distance of each speckle from the cellular center of mass for P-bodies and N protein. Bars show the mean with SEM and single cell values are displayed as dots for different time points post infection. **(C)** Analysis of per cell counts (abundance) and localization analysis for individual vRNA and viral mRNA species. ****, p≤ 0.0001; ***, p≤0.001; **, p= 0.001 to 0.01; *, p= 0.01 to 0.05. Significance was analyzed using one-way ANOVA and Tukeýs multiple comparisons test.

We also conducted additional qPCR experiments to validate our FISH data (Figure S9). For that purpose, we quantified vRNAs and viral mRNAs in cell pellets and cellular supernatants at different time points post infection. Overall qPCR data were highly consistent and strongly supported our microscopy results (Figure 3, Figure S9). For instance, both methods showed a higher abundance of vRNAs than mRNAs, but also generally higher levels of S-mRNA/vRNA than L-mRNA/vRNA (∼factor 10). In addition, our qPCR results indicate that mRNAs plateau after 120 h of infection, which we also observed by FISH for M-mRNA. In strong contrast, both qPCR and FISH show that vRNA levels further rise beyond 120 hpi.

### N protein is the dominant factor at late infection time points

Next, we performed analyses of all detected spots in concert, considering the frequency of individual vRNA and mRNA species, as well as P-bodies and N protein speckles (Figure 5). Our data indicate that the large majority (∼75%) of all identified puncta at 24 hpi were P-bodies, whereas N protein aggregates make up more than 50% after 10 days of infection (Figure 4A, B, C, Figure S10A). Furthermore, we monitored co-localization of all markers by calculating pairwise correlation coefficients (Figure 4D, Figure S10B). We found a robust correlation between S- and M-vRNA, with N protein and P-bodies at 24 hpi, whereas L-vRNA showed weaker, but significant correlation. After 10 days of infection almost all correlation levels increased markedly, except for vRNAs with viral mRNAs, where levels remained comparatively low. We further examined the frequency of different factors alone, or in complexes with each other (Figure 4C). For this purpose, different vRNA and viral mRNA species were not distinguished so that i.e., spots with either of the viral RNA segments score as vRNA positive. We observed that the frequency of individual P-body and individual vRNA spots decreased over time (black and cyan), while spots with N protein- vRNA (orange) and N protein-mRNA (pink) co-localization, as well as N protein only spots (blue) gradually increased (Figure 4C). Aggregates that contained all components, which could indicate active viral factories, were found to be rare at 24 h, peaked at 48 h and remained fairly low, but stable for the remainder of the time course (Figure 4C, striped pattern at the bottom). Importantly, the frequency of spots with N protein, vRNA and mRNA, but without P-body markers was very low (Figure 4C, grey bars), which supports the notion that viral mRNA only associates with other viral components in (P-body containing) viral factories and does not directly interact with vRNAs or N protein. Of note, a simulated dataset demonstrated that our analysis is highly specific and indicative of actual co-localization rather than random spot clustering as a result of high label densities (Figure S11).

**Figure 4:**
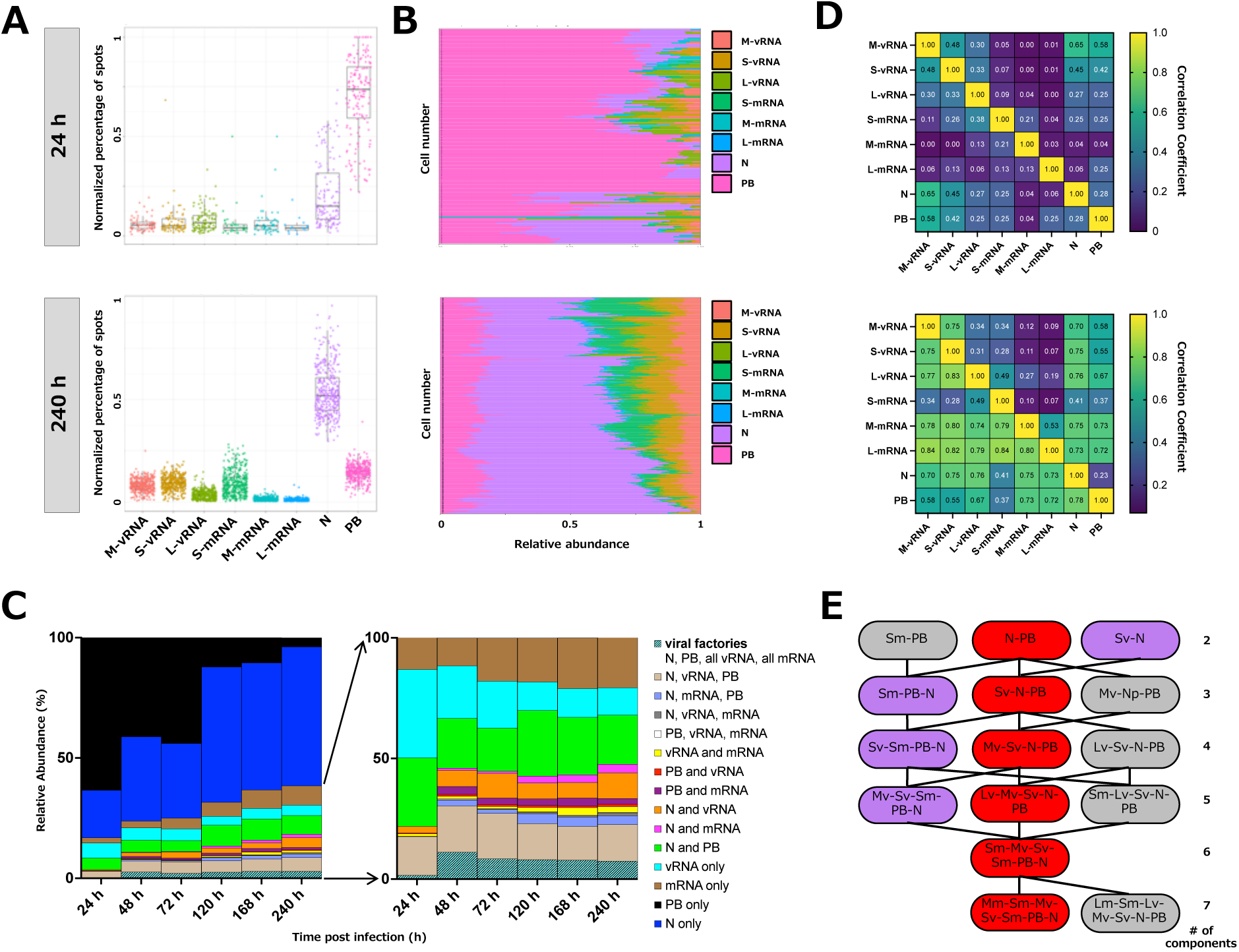
Quantitative analysis of spot ratios, measured by MuSeq-FISH. **(A)** Cell infected with PUUV (MOI=0.3) as shown in (Figure 3) were investigated by MuSeq-FISH and analyzed for spot ratios and co-localization of different markers. Mean per cell spot ratios at 24 and 240 hpi are shown in a Tukey boxplot representation. Upper bound of box, center and lower bound of the box represent the 75th percentile, the 50th percentile (median) and the 25th percentile, respectively. Upper and lower whiskers represent the maxima and minima of the boxplots showing the respective largest or smallest value within 1.5× interquartile range above the 75th or below the 25th percentile (bounds of box). Individual cell values are indicated by colored dots. (B) Relative spot ratio analysis for individual cells at 24 and 240 hpi. Each horizontal line reflects an individual cell, with colors indicating the respective spot species. The number of spots that were detected was counted and normalized to the total number of spots per cell. (C) Mean relative spot abundances in percent at different time points post infection. In the right panel N proteins and P-bodies are disregarded to highlight low abundance spot populations. (D) Pair-correlation of fluorescence signals at 24h and 240 h. (E) Potential assembly and clustering pathways in Puumalavirus infected cells. The most abundant multi-color complexes, with either 2, 3, 4, 5, 6 or 7 components are displayed in boxes, with lines indicating likely transitions to assemblies with higher complexity. Joining lines were drawn if boxes of adjacent rank (complexity) could be connected to the next through the addition of a single component. Sm=S-mRNA, Mm=M-mRNA, Lm=L-mRNA, Sv=S-vRNA, Mv=M-vRNA, Lv=L-vRNA, PB=P body, N=N protein.

**Figure 5:**
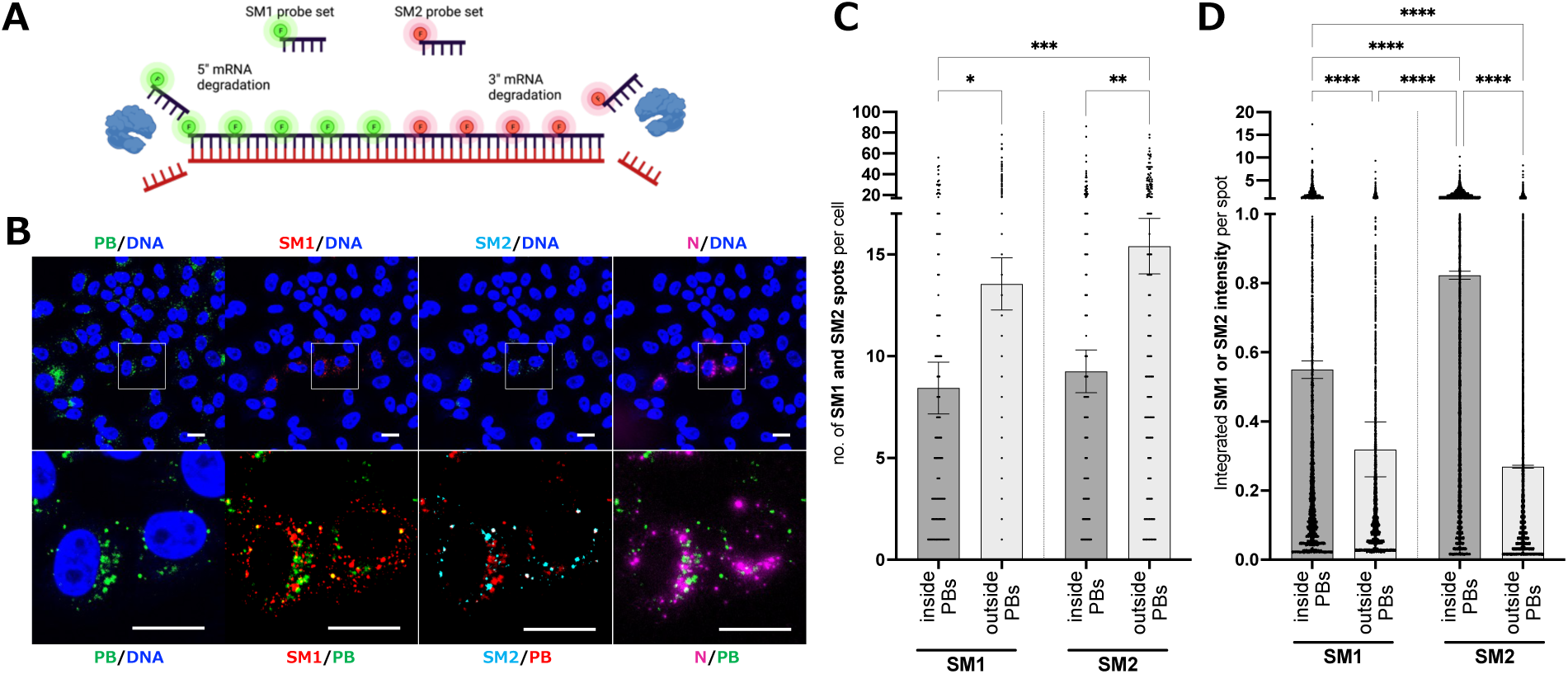
Site specific labeling of viral mRNA to study their degradation in P-bodies. **(A)** Schematic representation of the experimental setup for mRNA degradation analysis. **(B)** VeroE6 cells were infected with PUUV MOI 1 and stained at 72 hpi using anti-DCP1a (P-bodies) antibody, accompanied by FISH staining with S-mRNA probe subsets, SM1 and SM2, corresponding to the 5’ and 3’ ends of the S-mRNA. Representative fluorescence microscopy images, including counterstaining for DNA, PUUV N protein and P-bodies using DCP1a antibodies. Scale bars=10 µm. **(C)** Quantification of SM1 and SM2 speckle abundance and integrated signal intensity using CellProfiler. ****, p≤ 0.0001; ***, p≤0.001; **, p= 0.001 to 0.01; *, p= 0.01 to 0.05. All bars indicate the mean with SEM. Significance was analyzed using one-way ANOVA and Tukeýs multiple comparisons test. PB=P- body.

Ultimately, we sought to shed light on the orthohantavirus assembly progression and hierarchy. To that aim, we assessed the most frequently observed multi-marker spot with a certain level of complexity (single, double, triple, etc. positive) at each time point and utilized that information to build an assembly tree (Figure 4E). For instance, among the double positive spots, N and P-bodies was the most probable combination. For triple positive spots, N, P and S-vRNA was most abundant, and accordingly for high order complexes. This approach allowed us to propose an assembly order that starts with an association of N protein and P-bodies, which then incorporates different vRNA species and eventually also encompasses viral mRNAs (Figure 4E).

### PUUV mRNAs are preferentially degraded from the 5’ end in P-bodies

P-bodies have key functions in the degradation and catabolism of cellular transcripts (14). Different mechanisms exist that digest mRNA from either 3’ or 5’ ends, ultimately leading to the decay of P Body-associated mRNAs (15). To test for end-specific degradation of S-mRNA in host cell P-bodies we devised an approach, in which 5’ and 3’ ends of the viral mRNA were labeled with different FISH probes (Figure 5A), called SM1 and SM2, respectively. If S-mRNA would be preferentially degraded from either direction, we would expect to see differences in the SM1 over SM2 abundance in P-bodies.

We were able to detect both probes in infected samples (Figure 5B) and quantified the number of SM1 and SM2 speckles, as well as their integrated intensity for either P-body-associated or P-body-independent speckles (Figure 5C). Firstly, we found for both probe sets that significantly less S-mRNA speckles were associated with P-bodies then without (Figure 5C, grey and white bars). The overall number of SM1 speckles and SM2 speckles either in or outside the P-bodies however was identical for both probes arguing against a specific degradation direction. Interestingly, the integrated signal intensity of both probes was significantly higher in P-body positive than in P-body negative speckles, an observation that reflects a local enrichment of viral mRNAs in P-bodies (Figure 5D). Notably, SM1 seemed to have a significantly lower signal intensity in P-bodies than SM2, which could indicate a preferential degradation from the 5’ end of the transcript (Figure 5D, grey bars). This argument is supported by the fact that outside P-bodies, both SM1 and SM2 exhibited comparable fluorescence intensities. It is therefore unlikely that differences in the fluorescence properties of SM1 and SM2 labels account for lower signals of the P-body associated SM1- probes.

## DISCUSSION

In this study, we investigated and quantitatively assessed the spatiotemporal dynamics of orthohantavirus infection using multicolor FISH and IF staining. We comprehensively monitored the three orthohantavirus genomic vRNA segments and viral mRNAs throughout complete infection cycles, while concomitantly studying co-localization with the viral N protein, host actin, microtubules, and P-bodies (Figure 1).

### Co-localization of vRNAs with host factors and cellular remodeling upon infection

Previous studies have extensively investigated the role of cytoskeletal factors in orthohantavirus infection (5,7,11,16–18). At this point, it is well established that virus proteins colocalize with microtubules, actin and the intermediate filament protein vimentin (5,11,16,18). Moreover, it is known that a pharmacological disruption of the cytoskeleton leads to either a redistribution of viral proteins or, in many cases, to an inhibition of key steps of the virus replication cycle (7,11,16,17). A cellular remodeling has previously been shown upon Andes Virus (ANDV) infection, which markedly rearranges Vimentin (11), and for Tula Virus (TULV), where late/chronic stages of infection appear to recruit stress granules to filamentous N protein aggregates (5). In addition, we have recently shown that PUUV can induce a redistribution of filamentous actin in infected cells (7), an observation that was confirmed and significantly expanded in this study.

Our current experiments indicated a spatial correlation of filamentous actin with vRNAs and N protein, suggesting direct interactions of virus components with host cytoskeletal factors (Figure 1). Moreover, we found that P-bodies exhibit extensive co-localization with N protein and vRNAs, reflecting their crucial involvement in orthohantavirus replication (Figure 1D). However, we also observed a redistribution of P-bodies, microtubules and filamentous actin (Figure 1B), accompanied by an increase in the number of P-bodies upon PUUV infection (Figure 1C, Figure 3A). This finding points towards a marked, cellular remodeling by PUUV, which gradually intensifies throughout the infection cycle (graphically summarized in Figure 6). We additionally investigated P-body associated RNA degradation processes by utilizing a novel approach involving 3’ and 5’ vRNA probe sets. Our results suggest that S-mRNA degradation occurs preferentially from the 5’ end (Figure 5), however further, detailed studies are necessary to conclusively address degradation of virus mRNAs. Of note, a number of early studies has thoroughly investigated the role of P-bodies in orthohantavirus infection (3,4). Noteworthily, the crucial viral cap-snatching, taking place in these cellular entities, is not only a hallmark of the orthohantavirus genus, but a mechanism that is shared across the entire Bunyavirales order. We would like to point out that a remodeling of P-bodies, has not been previously reported, however a transient increase in the abundance of stress granules could be observed for both for PUUV and ANDV (19), which is reminiscent of changes in the P-bodies homeostasis we observed.

**Figure 6:**
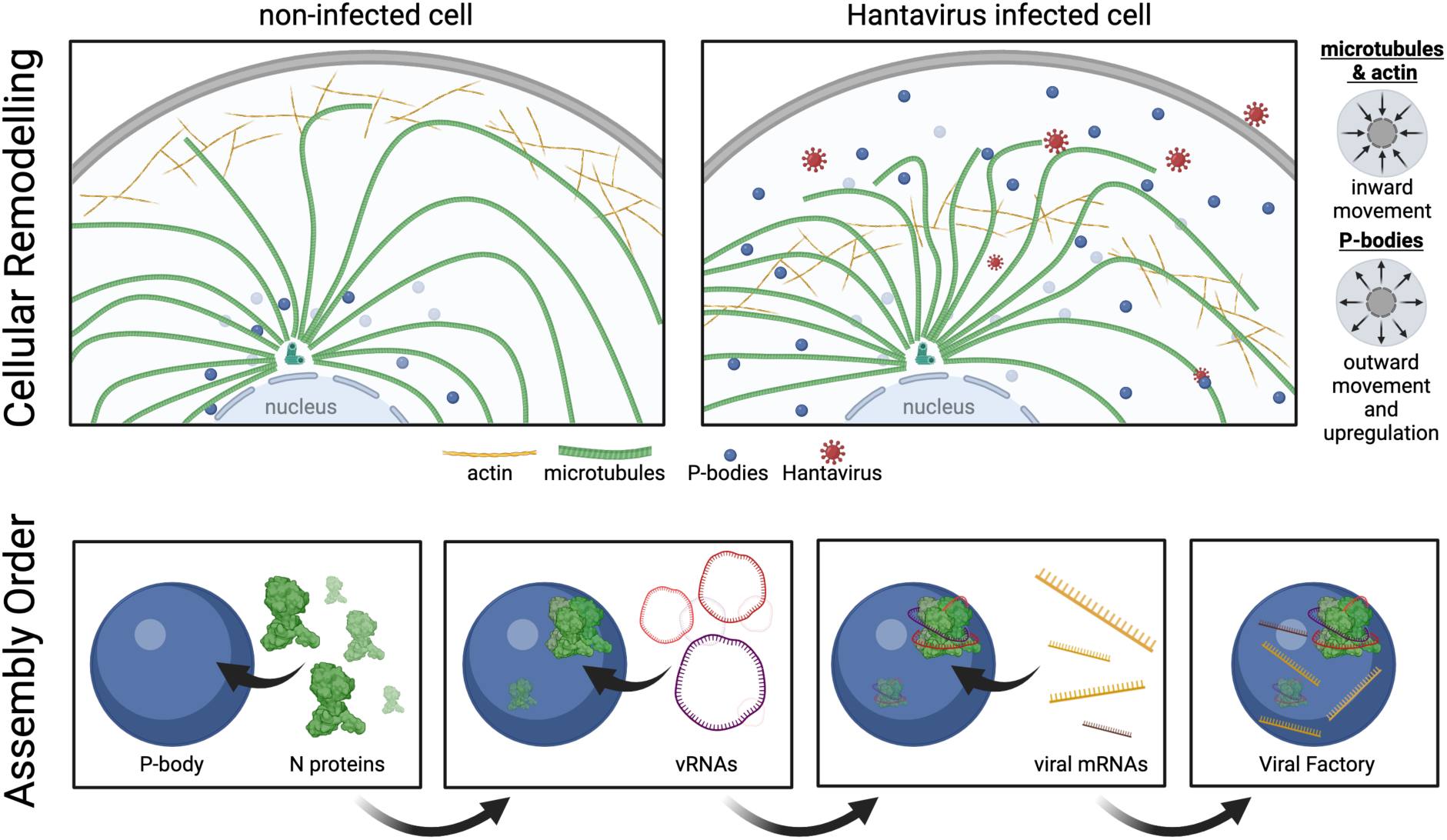
Graphical summary of cellular remodeling upon orthohantavirus infection and a putative orthohantavirus assembly regimen. Based on our experimental results we propose a model in which orthohantavirus infection induces a relocation of cellular actin and microtubules towards the cell center, whereas P-bodies exhibit an outwards movement and increase in overall abundance. Moreover, our data indicate that virus factories can form in infected cells through a stepwise association of N protein with P-bodies, followed by recruitment of v RNAs and eventually incorporation of viral transcripts.

At this point it is unknown what consequences the cellular remodelling has for the infected cells. We have previously reported that Orthohantaviruses exploit and even induce macropinocytic fluid-phase uptake, even prior to an infection. The actin remodelling we have described here and previously (7) could likely be involved in this manipulation of cellular functions. Interestingly, N protein has been reported to strongly co-localize with actin and P-bodies, even in absence of other viral components (18), thus one could speculate whether direct interactions between N and these cellular factors is contributing to their redistribution. It remains elusive however, how exactly cellular protein and mRNA homeostasis are affected by such virus-induced host cell rearrangements. Pinkham at al. have previously provided a comprehensive analysis of the transcriptional changes upon infection with Rift Valley fever virus, a related member of the Bunyaviridae family (20). The authors report a significant upregulation of several cellular transcripts, which could be a result of a P-body and therefore RNA-degradation dysfunction. Whether P-body remodelling and (putative) dysfunction are mechanistically connected, remains an open question.

### Spatiotemporal analysis of orthohantavirus replication and assembly

Another focus of our study was the interplay of virus genomic RNAs, transcripts and proteins in orthohantavirus infected cells. Throughout our FISH experiments we found discrete intracellular foci, with various combinations of vRNA and N protein (Figure 1, Figure 4), indicating the coexistence of multiple stages of virus assembly and replication within infected cells. Moreover, our intracellular localization analyses suggest disparities in the intracellular distribution between different virus mRNAs on one hand, and between virus mRNAs and corresponding vRNAs on the other hand (Figure 3). Of note, early infection time points exhibited a peak in vRNA-, but not mRNA-distances from the cell center, likely reflecting the virus inoculum (Figure 3). By analyzing the relative spot ratios, we observed a shift in the frequency of different spot species over time (Figure 4). In addition, our co-localization analysis revealed a marked correlations between S- and M-vRNA with N protein and P-bodies already at early time points, while co-localization of all markers clearly increased at later stages of infection (Figure 4). Overall, we found that N protein positive speckles dominated at late infection time points (Figure 4). Our quantification also revealed that P-bodies, over time, become heavily associated with virus components, with as few as 10 % not being associated with either vRNAs or N protein after 10 days of infection (Figure S9).

Based on our comprehensive MuSeq-datasets, we proposed an assembly pathway derived from the most frequently observed multi-marker spots (Figure 6). This pathway suggests an initial association of N protein with P-bodies, followed by the incorporation of different vRNA species and eventually virus mRNAs (Figure 6). Since virus mRNAs are not majorly incorporated into virus particles, the latter population of intracellular aggregates, positive for all markers, likely represent active viral factories, as previously described for bunyamwera viruses (21). Importantly, our assembly pathway indicates a successive recruitment of the different vRNA species, which progresses from the S-segment, over M- and then to the L-vRNA. This could suggest that vRNAs are added in the order of increasing complexity (Figure 6E).

However, it is important to point out that our study did not attempt to provide a quantitative, molecular analysis of virus particles or intracellular assembly sites. Rather than assessing the composition of such entities on a single molecule level, our goal was to visualize and quantify the interplay between different virus components, but also their association with cellular factors, such as P-bodies or cytoskeletal proteins. Previous studies thoroughly investigated and described the packaging and intraviral organization of different bunyaviruses (22–25), which will likely translate, at least to some extent, also into the orthohantavirus genus. Nonetheless, additional studies are necessary to reveal whether packaging dynamics and kinetics are shared throughout the Bunyavirus order or differ for specific species.

Overall, our study provides comprehensive insights into the spatial organization and interactions of virus nucleic acids, N protein, and host cellular components upon infection of an emerging human pathogenic member of the orthohantavirus genus, thereby unraveling the intricacies of orthohantavirus replication and assembly mechanisms, and emphasizing the pivotal role played by the cytoskeleton and P-bodies in the virus life cycle.

## ACKNOWLEDGMENT

A.H. was supported by CRC 1449 “Dynamic Hydrogels at Biointerfaces” (DFG, German Research Foundation), Project ID 431232613–SFB 1449.

## AUTHOR CONTRIBUTIONS

HSS, AP, MS, NN, SW, LCA carried out the experiments. MJL, PTW, DB, DHK, AH, RS supervised the research. HSS, AP, MS, KP and RS analyzed the data. HSS, AP, DHK and AH prepared the manuscript.

## ABBREVIATIONS

IF: immunofluorescence
HCPS: hantavirus cardiopulmonary syndrome
HFRS: hemorrhagic fever with renal syndrome
MCC: multi-segment complex
(MuSeq-)FISH: (multiple-sequential) fluorescence *in situ* hybridization
N: nucleocapsid
P-bodies: processing bodies
PUUV: Puumala virus
RdRp: RNA- dependent RNA Polymerase polymerase
v(m)RNA: viral (messenger) RNA
vRNP: viral ribonucleoprotein complex

## SUPPORTING INFORMATION CAPTIONS

**Figure S1:** (A) Confocal images of VeroE6 cells subjected to IF staining and L-vRNA FISH at 72 hpi. The upper left panel shows mock-infected (uninfected), the lower panel infected samples (MOI=1). Colored labels indicate the displayed target molecule. The boxed area in the center overlay image is shown magnified on the right. Thin dashed lines in the magnification image indicate regions of interest used for line plots in C. (B) S-vRNA FISH and M-vRNA FISH in infected cells stained by IF for N protein. All images show maximum intensity projections of z-stacks, obtained by confocal microscopy. DAPI was used as a DNA counter staining (blue). All scale bars correspond to 10 µm. (C). Line plots, showing signals in the vRNA (red) and N protein (green) channels.

**Figure S2:** Viral RNA associates with actin accumulations, microtubules and P-bodies. (A) Combination of TRITC-phalloidin (actin), (B) anti-α-tubulin (microtubules), (D and E) anti- DCP1a (P-bodies) and N proteins staining with FISH in VeroE6 cells infected with PUUV (MOI=1) for 72 h. Images show equatorial slices obtained by confocal microscopy. Hoechst 33342 was used as a DNA counter staining (blue). Scale bars=10 µm. (C and F) Line plot analysis of fluorescence microscopy images. Plots show normalized intensities in different fluorescence channels (blue, red and green), along manually selected, linear regions of interest (dashed lines in fluorescence images). Analysis of cells stained for either (C) α-actin or tubulin, as well as (F) P-bodies, along with N protein, DNA and S-vRNA. All line plots indicate regions with peak vRNA/N protein intensities that coincide with local actin, α-tubulin or P-body maxima. Additional examples images of all three vRNA segments can be found in Figure S3.

**Figure S3:** vRNA co-colocalization with cellular markers (related to Figure S2). Additional examples for stainings shown in Figure 2. (A) Immunofluorescence staining with anti-tubulin (microtubules) and anti N protein antibodies in combination with FISH labeling of S-vRNA, (B) M-vRNA, or (C) L-vRNA. (D) Anti-DCP1a (P bodies), N protein and FISH staining for S-vRNA in VeroE6 cells infected with PUUV for 72 h. (D, E, F) Additional examples of a DCP1a, N protein and FISH-staining for S-vRNA, M-vRNA and L-vRNA, respectively. All images show equatorial slices obtained by confocal microscopy. Hoechst 33342 was used as a DNA counter staining (blue). Scale bars=10 µm.

**Figure S4:** Image segmentation and analysis. (A) Schematic representation of our custom-made image segmentation pipeline. Briefly, nuclei are identified in DNA counter stainings, followed by recognition of the cytoplasm in IF staining’s of abundant cellular markers or in transmission light images. Finally, P-body, N protein or vRNA puncta were identified independently. (B) Magnification of representative spot detections conducted by our image segmentation pipeline. (C) Schematic representation of a radial distribution analysis, which quantifies fluorescence intensities as a function of their distance from the center of individual cells. Briefly, the analysis first defines radial bins, then sums the overall intensities from equivalent bins of multiple cells and finally displays them in line graphs. (D) Schematic representation of co-localization analysis. Each identified spot is analyzed for co-localization of different fluorescence signals and summed up on a per cell basis. Frequencies of different spot species (single positive, double positive, triple positive) are then calculated for all analyzed cells and displayed in pie charts.

**Figure S5:** Radial distribution analysis for DNA stainings in infected and non-infected cells. Mean fractional intensity was calculated using the CellProfiler module MeasureObjectIntensityDistribution, binning the cell body into 10 section starting from the center of the nucleus going outwards. Several dozen to several hundred cells from different fields of view were analyzed for each plot. Error bars show the standard error of the mean (SEM).

**Figure S6:** Cellular remodeling of human pulmonary microvascular endothelial cells. Cells were infected with PUUV (MOI 0.5) and subjected to immunofluorescence staining at 72 hpi. Then, samples were imaged by confocal microscopy and analyzed using CellProfiler. (A) Representative fluorescence microscopy images. Marker colors are indicated above and below the micrographs. Scale bars=10 µm. (B) Quantitative image analysis using Cell Profiler. Left: integrated Actin and PB intensities were assessed in both infected and non-infected samples on a per-cell basis. Each dot represents individual cells and error bars show SEM. Significance was assessed by unpaired Student’s t test. Center and right: Mean fractional intensity (MfrI) was calculated using the Cell Profiler module MeasureObjectIntensityDistribution, binning the cell body into 10 sections starting from the center of the nucleus going outwards. Several dozen to several hundred cells from different fields of view were analyzed for each plot. Error bars show the standard error of the mean (SEM), and statistical significance was assessed by comparing PUUV and non-infected samples at each bin.

**Figure S7:** MuSeq-FISH working principle. (1) Cells are infected with Orthohantaviruses and subjected to (2) FISH staining, followed by (3) confocal fluorescence microscopy and (4) a Formamide wash that removes all FISH probes. Subsequently, another round of FISH staining can be conducted.

**Figure S8:** Imaging of PUUV vRNAs and their transcripts in infected cells. VeroE6 cells were infected with PUUV for 240 h and analyzed using MuSeq-FISH. (A) Overview and overlay images of a representative region of interest, showing each staining individually: S-mRNA and S-vRNA in green, M-mRNA and M-vRNA in red, L-mRNA and L-vRNA in cyan, N protein in yellow, P-bodies (PB) in purple, DNA in blue and cell mask staining in grey. Overlays display stained factors as indicated below. (B) Magnification of the boxed region shown in the cell mask image in (A). Scale bars=10 µm. Images represent maximum intensity z projections.

**Figure S9:** Quantification and kinetic analysis of all PUUV viral mRNA and vRNA species at different time points post infection using qRT-PCR. Vero E6 cells were infected with PUUV MOI 0.3. Cells and supernatant were harvest at the indicated time points. Both graphs show the mean RNA levels with the SEM. The samples were done in triplicates.

**Figure S10:** Quantitative analysis of spot ratios as measure by MuSeq-FISH. (A) Mean per cell spot ratios and relative spot ratio for all time points post infections are shown. Per cell spot ratios are displayed in a Tukey boxplot representation. Upper bound of box, center and lower bound of the box represent the 75th percentile, the 50th percentile (median) and the 25th percentile, respectively. Upper and lower whiskers represent the maxima and minima of the boxplots showing the respective largest or smallest value within 1.5× interquartile range above the 75th or below the 25th percentile (bounds of box). Individual cell values are indicated by colored dots. Relative spot ratios are shown in horizontal lines, reflecting an individual cell, with colors indicating the respective spot species. The number of spots that were detected was counted and normalized to the total number of spots per cell. (B) Pair-correlation of fluorescence signals at 24h and 240h. (C) Relative abundance of different spot species at different time points post infection. Individual pies indicate the fraction of a certain spot species (see legend).

**Figure S11:** Co-localization analysis of a simulated data set and a MuSeq-FISH data set of PUUV infected Vero E6 cells. To ensure accurate MSC grouping and proper assessment of colocalization data, an artificial data set was created. This data set consisted of a simplified cell shape model with a randomized spot distribution. The spot densities chosen for the simulated cells matched the spot densities previously measured in PUUV infected cells. All segments were equally represented in the simulated cells or had a weighted occurrence. Higher spot densities resulted in higher colocalization ranks. Only at a maximum spot density of 0.5 spots/µm3 (equivalent to approximately 750 spots per cell) and with an equal probability of the individual segments in a cell, were higher MSC ranks up to 3 determined with a likelihood of about 12 % (A). The rate of random colocalization in the weighted occurrence of the individual segments was always below 10%, primarily due to the clustering of PB and N. Contrary to the simulation, PUUV infected cells showed higher co-localization ranks up to 8 (B), despite having fewer spots compared to the maximum in the simulated data set (Figure 7A). Therefore, higher colocalization ranks observed in PUUV infected cells are unlikely to be a result of random co-localization caused by high spot densities.

